# Impact of resistance on therapeutic design: a Moran model of cancer growth

**DOI:** 10.1101/2023.08.28.555214

**Authors:** Mason S. Lacy, Adrianne L. Jenner

**Affiliations:** School of Mathematical Sciences, Queensland University of Technology, Brisbane, QLD, Australia

## Abstract

Resistance of cancers to treatments, such as chemotherapy, largely arise due to cell mutations. These mutations allow cells to resist apoptosis and inevitably lead to recurrence and often progression to more aggressive cancer forms. Sustained-low dose therapies are being considered as an alternative over maximum tolerated dose treatments, whereby a smaller drug dosage is given over a longer period of time. However, understanding the impact that the presence of treatment-resistant clones may have on these new treatment modalities is crucial to validating them as a therapeutic avenue. In this study, a Moran process is used to capture stochastic mutations arising in cancer cells, inferring treatment resistance. The model is used to predict the probability of cancer recurrence given varying treatment modalities. The simulations predict that sustained-low dose therapies would be virtually ineffective for a cancer with a non-negligible probability of developing a sub-clone with resistance tendencies. Furthermore, calibrating the model to *in vivo* measurements for breast cancer treatment with Herceptin, the model suggests that standard treatment regimens are ineffective in this mouse model. Using a simple Moran model, it is possible to explore the likelihood of treatment success given a non-negligible probability of treatment resistant mutations and suggest more robust therapeutic schedules.

## 1. Introduction

Resistance of cancer cells to chemotherapy is the first cause of cancer associated death and overcoming resistance is a critical focus for many researchers [1]–[4]. Resistance to cancer therapeutics is multifaceted and can arise due to several factors [5]. Some cancers are intrinsically resistant to treatments due to genetic complexity [6] and other cancer types can develop resistance during treatment [7], [8]. Drugs can be selected to target a specific cancer type, which presents a lower probability of developing a resistance to that drug [9]. Overall, more research into the impact of treatment resistance is desperately needed to improve our ability to overcome or better treat cancer patients.

Resistance to targeted therapies can arise from selective growth of pre-existing subclones within the bulk of the tumour that carry drug-resistance mutations, and thus have a survival advantage [7] (**Figure 1**). For example, some breast cancer cells will overexpress the HER2 protein on the cell surface. This overexpression is due to a driver mutation that accumulates over time [10], [11]. A drug known as Herceptin is able to target cells overexpressing HER2 and induce death [12]. Unfortunately, a significant number of patients do not benefit from this therapy, highlighting the need to understand the mechanisms of cancer resistance [13] and the impact of therapeutic protocol design.

**Figure 1.**
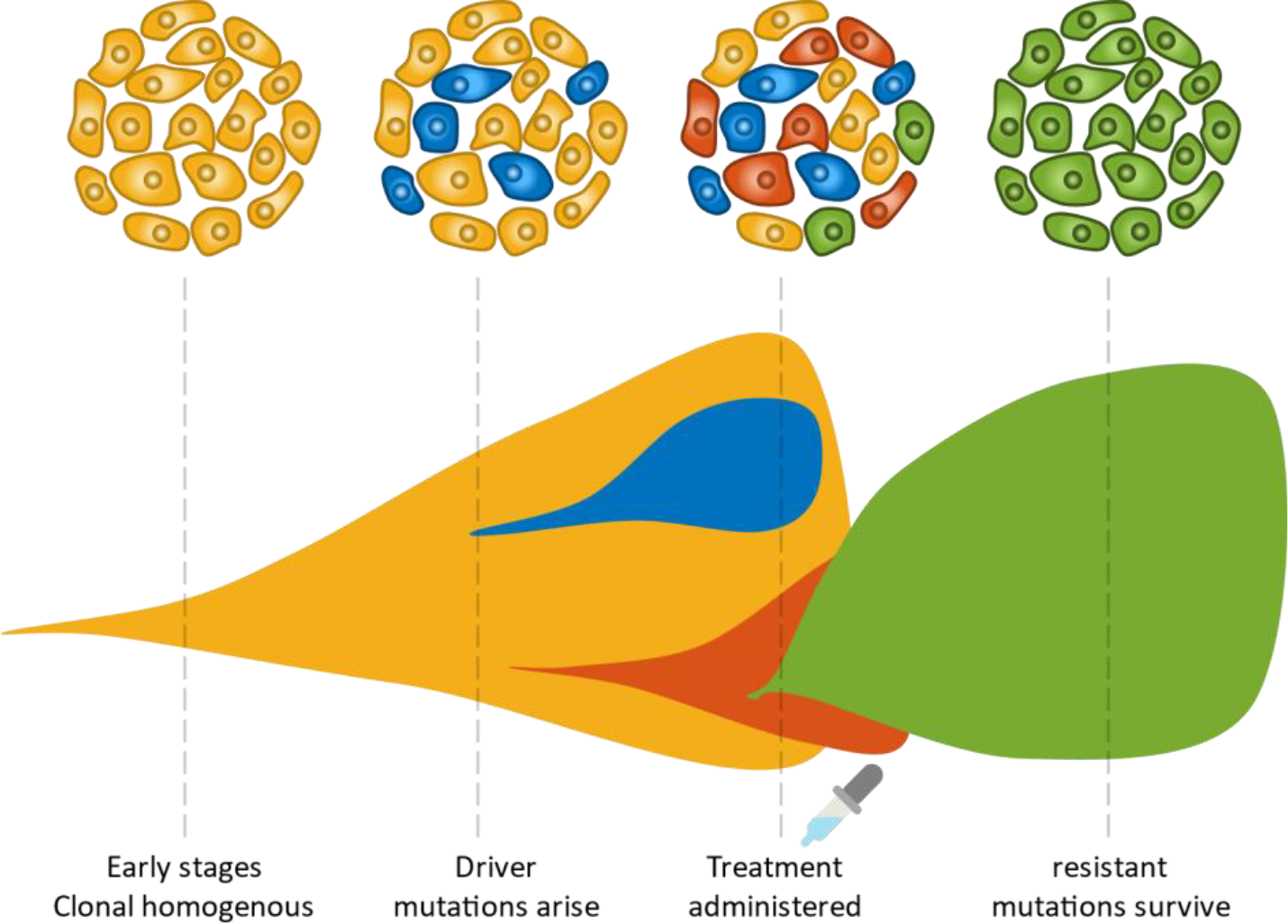
Schematic depicting tumour evolution over time and the fixation of a resistant clone following therapy. As a cancer grows, it is subjected to various pressures which can cause mutations to arise. Some clones may contain mutations that may be more adept at coping with treatment or provide a fitness advantage to those cells, such as faster proliferation rates, and we denote these as driver mutations. After treatment, often cells with driver mutations conferring resistance and/or fitness advantages will expand in number. In some cases, this can result in a tumour that is no longer as genetically complex. Most importantly, these tumours are often no-longer sensitive to the original therapy.

Standard clinical protocols for cancer treatment, such as chemotherapy, typically employ the maximal drug dose that can be tolerated by the patient, often referred to as the Maximum Tolerated Dose (MTD) [14]. The administration of chemotherapy at reduced doses given at regular, frequent time intervals, termed ‘metronomic’ chemotherapy, presents an alternative to standard MTD [14]. In a similar manner, adaptive therapy is an approach to treatment based on maintaining a proportion of a treatment-sensitive population and including treatment holidays, where no drug is administered [15], [16]. Often, a single systemic administration (or infusion intravenously) can result in significant systemic toxicity and only a fraction of the injected dose reaching the tumour. As such, there has been a growing interest in the development of localised targeted delivery systems which can modify the biodistribution of drugs [17]– [19]. These include biodegradable implants [20] or nanocarriers [21]. Often these sustained-release systems provide a lower drug dosage over a longer period, however, not much is known about their success in the presence of resistant mutations [22], [23]. More research into minimising the effects of drug resistance is required, highlighting the need to implement and test novel methods in simulated environments.

Unsurprisingly, cancer growth is an extremely heterogeneous disease, both in the way it presents across the human population, and in the inherent cellular subclones of a tumour (**Figure 1**). Cancers are usually comprised of multiple cell types, often referred to as clones, that can differ by the passenger and driver mutations they contain. Clones can give rise to specific cell lineages and the variation within a tumour is referred to as intra-clonal heterogeneity [24]. This presence of clones in a cancer cell population results in behaviour similar to that of Darwin’s evolution theory, which can promote drug resistance as clones compete for dominance [23]. Treatment can also facilitate this behaviour by causing the death of some cancer clones which could be benign, and supporting the more rapid growth of resistant and aggressive clones [25], so care must be taken during therapy.

Mathematical modelling has been used for many years to capture cancer growth and treatment resistance [26], [27]. While it is possible to use deterministic models to capture this physical system [28], [29], stochastic mathematical modelling provides a way to more accurately account for the inherent randomness present in cancer growth and treatment [30], [31]. Some of the stochastic techniques used to model cancer resistance include branching processes [32], [33], birth-death processes [34], [35], Wright-Fisher models and agent-based models [36]–[38]. A Moran process is a stochastic model that considers a fixed population of size *N* with a fixed number of states within the population. Transitions between states in each time step is governed by some probability. Moran models are often used for modelling cancer growth, which is driven predominantly by cell division and death [39]–[41], and are able to capture the interactions between healthy and cancerous cell clones using different fitness advantages *f* [27].

In evolution, cells or animals are considered to hold a fitness advantage if some inherent characteristics provide them with a benefit. For example, cancer cells may be considered more fit if they are a product of a driver mutation that provides them with an ability to proliferate more rapidly. In Moran models, cell fitness affects the probability of a particular cell being selected to reproduce, which could lead to fixation of a more fit cell type. West *et al*. [39] present a Moran model that considers multiple clones of both healthy and cancerous cells. Here, passenger mutations provide transitions between different healthy cell states, where some of these healthy cells have a higher probability of providing a driver mutation to cancerous cells. They also consider applying concepts seen in the prisoner’s dilemma evolutionary game to select the fitness for healthy and cancerous cells. They identify that treatment should be administered in the early stages of the cancers growth to prevent genetic complexity. Similarly, Heyde et al. [40] finds that the Moran process accurately describes the driver mutations found in hematopoietic stem cells.

Moran models are also used to track the probability of tunneling, which occurs when a cancer mutant reaches fixation before the original cancer. Komarova [42], Haeno et al. [43] and Jackson et al. [44] each considered a Moran model with three cell types, where the third type is a mutation from the original cancer with higher fitness. Each of these models discover that tunneling occurs naturally at some rate depending on the fitnesses of the cancerous and mutated cells. Similar Moran models have been applied to create more complex stochastic models that consider spatial information about the cells, like ‘warlock’ [45], or the one-dimensional model presented by Komarova [42]. This one-dimension spatial Moran process models the generation of cancer mutants more accurately than the non-spatial model, which seem to underestimate these mutations. Werner *et al*. [46] used a two type Moran model to capture wild type leukemic cancer cells sensitive to Imatinib and Imatinib resistant cancer cells. They compared the dynamics of their model to *in vitro* experiments to determine the fitness of different cell populations. Altrock and Traulsen [47] determine analytical expressions for the fixation probabilities of a two cell Moran model.

Moran modelling to date has been insightful for standard dosing regimens. In this work, we investigate the effects of resistance in MTD versus sustained low-dose treatment. To do this, we develop a simple Moran model to capture the development of chemotherapy resistance during cancer growth and treatment. In section 2, we detail the modelling assumptions and experimental data used in this work. In section 3, we investigate the effects of different treatment modalities and discuss the implications of our findings for future cancer treatments.

## 2. Methods

### 2.1. Moran birth-death process

The Moran process is a Markov process that models stochastic dynamics of a population with constant size *N*. In this work, we present a Moran model that describes the interactions between healthy cells and different clones of cancerous cells (**Figure 2**). Considering in the initial stages of cancer growth there are two cell types, cancerous (type *c*) and healthy (type *h*), with corresponding finesses *f*_*c*_ and *f*_*h*_. The populations of each cell type (*N*_*c*_ and *N*_*h*_ respectively) can be tracked in any given simulation, where the total population of cells remains constant at *N* = *N*_*c*_ + *N*_*h*_. In a single timestep of this two-cell Moran model, there are three possible outcomes (i) cell type *c* proliferates and cell type *h* dies, (ii) cell type *h* proliferates and cell type *c* dies, or (iii) cell type *c* and *h* undergo no change. Each option has an associated probability:

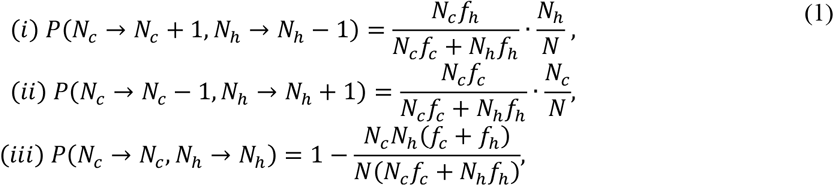

where the probability of each transition is proportional to the probability of a cell of a particular type being chosen to reproduce and a cell of a particular type being chosen to die. Specifically, we assume the probability of being selected to die is proportional to the relative size of the cell’s type in the total population *N*. In contrast, the probability of a cell being chosen to reproduce is proportional to its fitness *f*. Formulating the probabilities in this manner is similar to previous work by West *et al*. [39], Altrock *et al*. [27], and many others [42]–[44].

**Figure 2.**
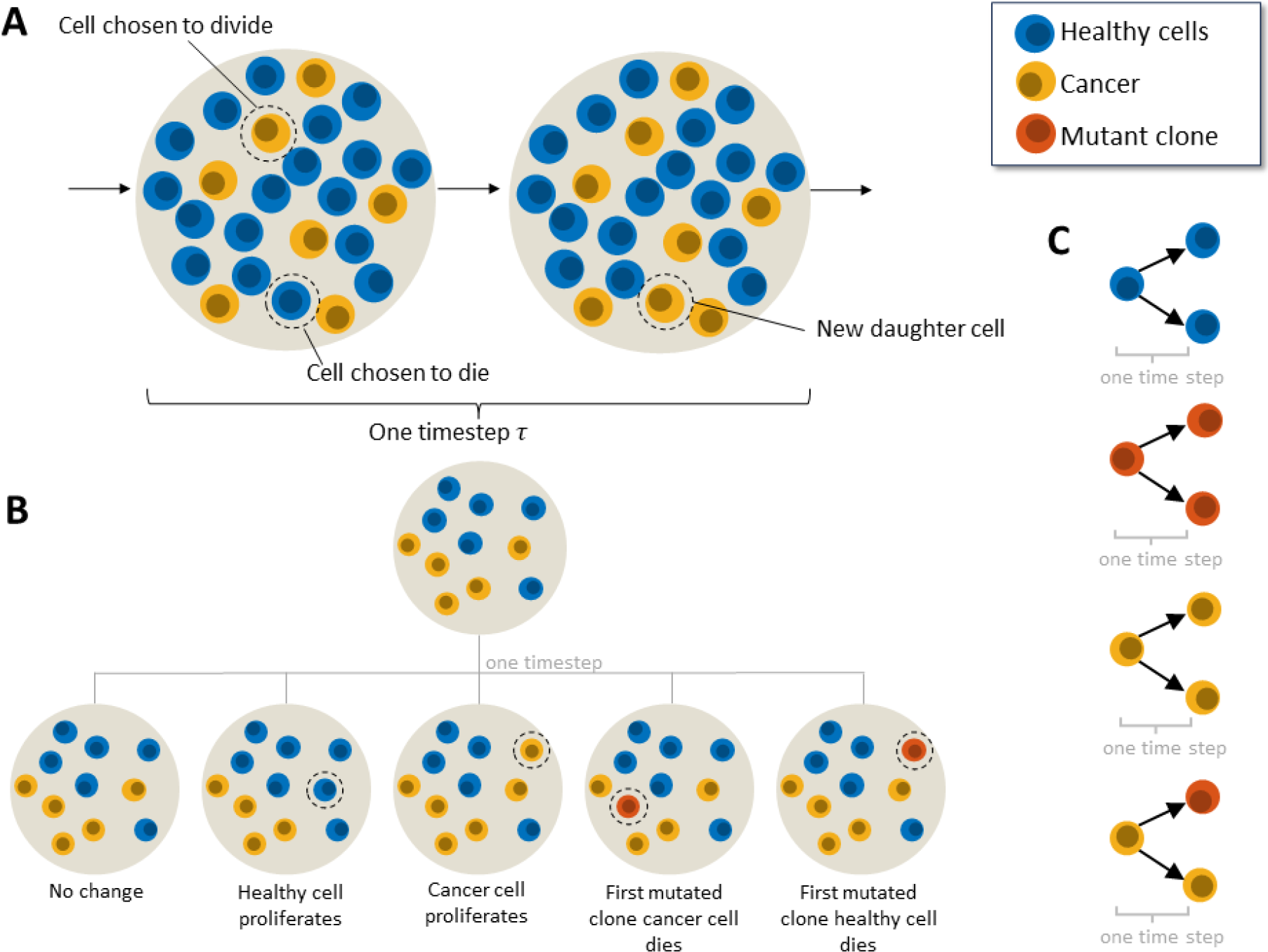
Moran model for cancer cell growth and mutation. (A) We assume initially there are two populations of cells: healthy and cancerous. In a single timestep τ one cell is chosen to divide and one cell is chosen to die based on the probabilities given in **Eq. (1)**. The cell chosen to die is replaced with a cell of the type chosen to proliferate. (B) Including mutations into the model (red cell) initially gives rise to 5 possible outcomes in a single time step where only healthy and cancerous (non-mutated) reside in the population. Either there is no change in the population, due to a cell of the same type being chosen to divide and die, a cancer cell is chosen to proliferate, and a healthy cell is chosen to die so the cancer population grows, a mutated clone is created and replaces a healthy cell, or a mutated clone is created and replaces one of the original cancer cells. (C) Lineage rules for each individual cell type. Healthy cells can divide into healthy cells, cancer cells can divide into cancer cells or mutant cells, and mutant cells can divide into mutant cells. Note that, for division to occur, a cell of a different type must be selected to die.

Of note, if the fitnesses are equal (i.e. *f*_*c*_ = *f*_*h*_), then (i) and (ii) are equal probabilities and it is with equal probability that cell type *c* or cell type *h* increases in a single timestep. Here, the probability of being selected to proliferate is proportional to the size of that cell’s type in the population, similar to the probability of being selected for removal from the population. Given this, and the fact there is a fixed total cell count *N*, we expect one cell type to fixate eventually. While the law of large numbers tells us that the average would be 50% of each cell type and we see this when we average many simulations, the cell populations won’t necessarily remain at this value given that once they reach *N* or 0 they are fixed there for all cell divisions when mutation is not considered (**Figure 3** and **Figure S1**).

**Figure 3.**
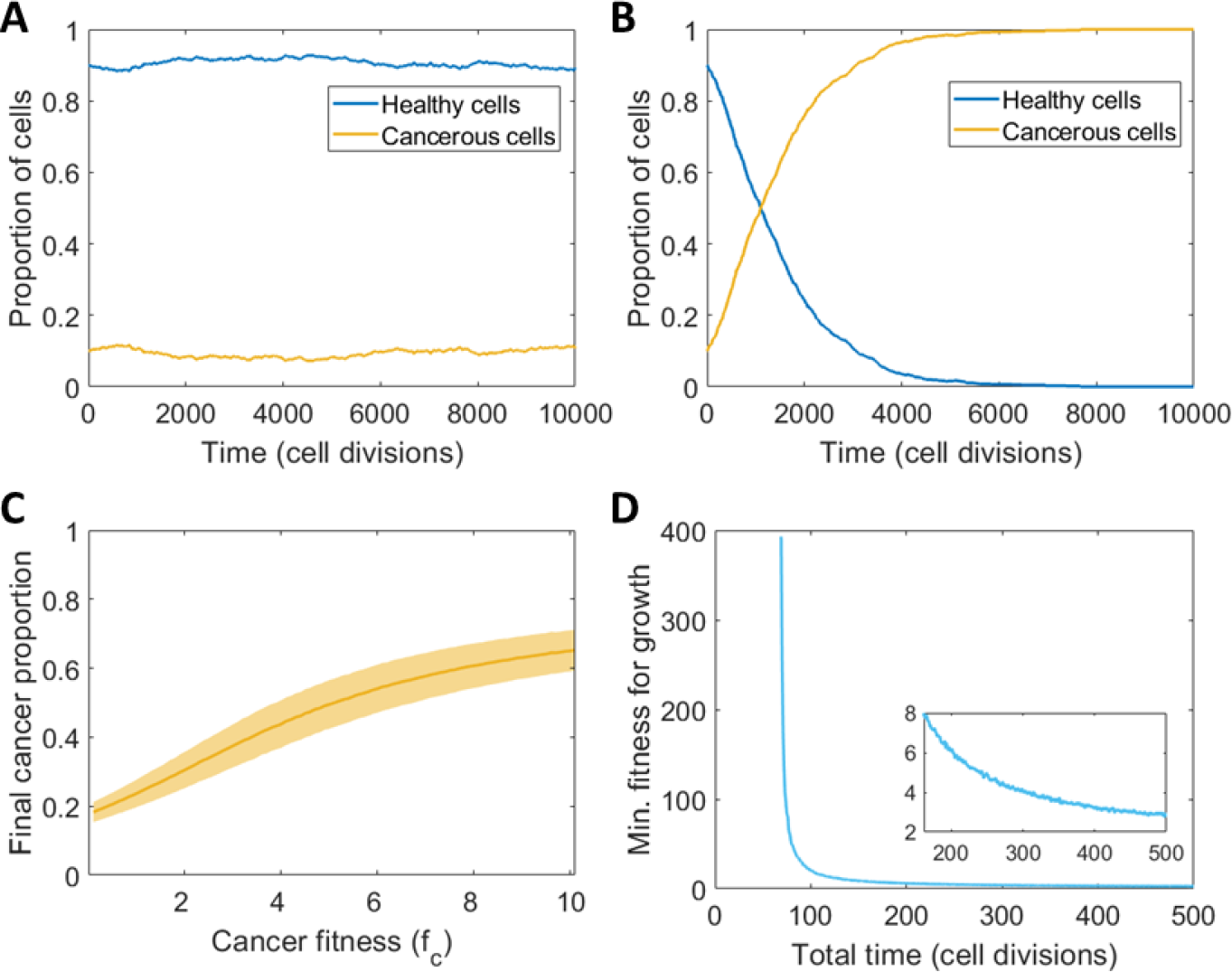
The predicted dynamics of the cancer cell-healthy cell Moran model in the absence of mutation and treatment. (A, B) Typical simulations of the Moran model for (A) equal fitness (*f*_*c*_ = *f*_*h*_ = 1) and (B) unequal fitness (*f*_*c*_ = 5, *f*_*h*_ = 1) with standard deviation of *n*_*total*_ = 100 simulations in **Figure S1**. (C) Simulation of the Moran model for total cell divisions *M* = 100 to find the final cancer proportion for different *f*_*c*_, average and standard deviation of *n*_*total*_ = 1000. (D) Required minimum cancer fitness to reach over 50% of the population being cancerous, i.e. *N*_*c*_ > 0.5*N*, where *N* = 100 for varying total time, i.e. number of cell divisions *M*.

### 2.2. Modelling mutation and intra-clonal heterogeneity

Using this framework, mutations from the original cancer into cancer clones are considered, represented by a fixed chance that a clone will emerge when the original cancer reproduces. This will be modelled with one cancer clone to represent the growth’s genetic complexity. Our simulation does not consider back-mutation, only driver mutations towards more fit cancer types. Passenger mutations are not present in this model as equally fit clones of healthy or cancerous cells are not considered. Rather, each cell type is considered as the collection of all cells with the same fitness advantage, regardless of potential genetic differences.

For this model, mutation introduces the type *m* cancer clone, referred to as mutated cancer, with corresponding mutation probability *r*_*m*_ (**Figure 2B**). The chance that a mutation occurs when type *c* cells reproduce, i.e. the probability of mutation, is given by

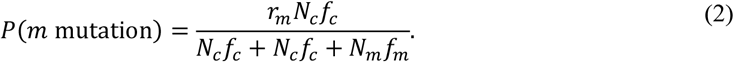

This could be extended for mutations of type *m*_2,_ … *m*_*n*_ cells equivalently, where their mutations can occur alongside each other. After the first driver mutation to a mutated cell type occurs, this mutation continues and their reproduction and death occur in the same way as type *c* and *h* cells, with fitness *f*_*m*_.

### 2.3. Modelling treatment

Treatment can be represented in the model as a reduction of cancer fitness; either reducing their reproductive advantage over healthy cells or introducing a reproductive disadvantage. Depending on the treatment type, some drugs can be more or less effective on different cancer clones [9]. This is equivalent to considering a non-uniform effect of treatment on the fitness values for each clone. Typically, treatment may be targeted towards the original cancer, but it could also be targeted towards a mutant clone [9].

In the model, treatment is introduced by considering some efficacy for different cancer clones, such as eff_*c*_ and eff_*m*_, which describes how effective treatment is for those cell types (0 having no effect and 1 being completely effective). Using these, the fitnesses of the cancer clone cells are reduced while treatment is active to give a modified fitness:

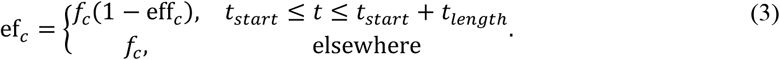

Here, treatment is only assumed to affect the reproduction of cancerous cells, and not increase the chance of death. Specifically, a single short high dosage treatment may be considered, where the treatment efficacy remains constant within some period with a length of *t*_*length*_ starting at *t*_*start*_, and treatment is not active outside of the period. It may also be the case that a lower dose is injected over a larger period, which considers a lower efficacy.

Given that a drug’s presence in the body is expected to reduce after it is administered, treatment effectiveness could also reduce over time. We assumed a drug’s half-life *t*_1/2_ may be used to model a waning treatment efficacy which is function of time since administration. Thus, an alternative model to **Eq. (3)** for treatment captures the drug efficacy decreasing over time by a decaying exponential, where it begins at the typical efficacy at the start of treatment and decays over time, as follows:

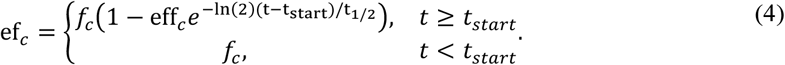

See **Eqs. (6)-(12)** in the **Appendix** for a full list of transitional probabilities including those under the effect of treatment.

### 2.4. Analytical expressions of the Moran model

Although simulations of the Moran model provide an understanding of the random nature of this process, it can sometimes be more beneficial to use an analytic expression for the expected results. Analytic expressions can reduce the computational cost previously incurred by evaluating *n*_*total*_ simulations to achieve the average or expected result. Using the probabilities governing cell reproduction and death in **Eq. (1)** (i-iii), the expected evolution of the cancerous cells (type *c*) over cell divisions (*n*) can be determined by the recursive sequence.

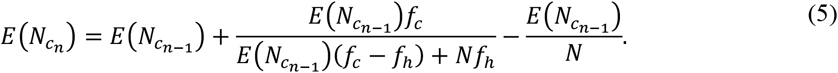

given some initial cell count 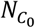. This definition holds similarly for the expected evolution of healthy cells (type *h*). This model closely matches the average Moran simulation when the initial number of cells is higher than ∼10% of *N*. Below this threshold, the average simulation is affected by trajectories where the cancerous cell type dies out due to stochastic fluctuations. This produces an average growth that is not expected of a typical simulation and does not match **Eq. (5)**. Mutation can be introduced into **Eq. (5)** by applying the mutation probability (**Eq. (2)**) to develop a recursive model for the expected evolution of mutated cancer cells (type *m*). Treatment can be introduced by allowing fitness to be varied as in **Eq. (3)** with constant efficacy, or **Eq. (4)** with treatment efficacy decreasing over time.

### 2.5. Experimental measurements of tumour growth

In this work, three sets of tumour growth measurements were used to validate and calibrate the model. The first measurements used were control tumour growth measurements form Laird [48], [49] who measured the growth of different cell lines in a rat tumour model. They normalised the time unit for 19 different tumour growth data sets using each data sets doubling time, this way all individual cancer line measurements were on the same normalised time axis. The efficacy of Herceptin was measured by Lewis Phillips *et al*. using an *in vivo* human breast cancer mouse model [12]. Tumour volume was measured with and without administration of Herceptin. Injections were given once every 3 weeks for three total injections.

### 2.6. Numerical simulations of the model and parameter estimation

Numerical simulations of the Moran model are performed using MATLAB (R2023a). When estimating parameters from data, the analytic recursive model in **Eq. (5)** is used for consistency and computational efficiency to estimate parameters. The Moran model is then simulated with the obtained parameter estimates and the mean of 100 simulations is confirmed to match the data. In cases where the initial number of cells are not relatively low, the average of 100 simulations matches the expected result closely within the data range. In both case studies considered in Section 3.4, the ratio of fitness between cancerous and healthy cells (i.e. *f*_*c*_ /*f*_*h*_ or *f*_*m*_/*f*_*h*_) is fit to data, alongside other measures such as mutation rate *r*_*m*_, treatment efficacies and measurement unit scaling. In the first case, the number of cell divisions per unit along the data’s horizontal axis is fitted to data, and when fitting treatment data, the efficacy of treatment against the original cancerous cell type is investigated. The complete expected Moran simulation is calculated using the recursive model for different combinations of these parameters and the Root Mean Square Error (RMSE) between this result and the data is recorded. For each complete simulation, the parameters that gave this error is recorded if the error is lower than the error given by the previous optimal parameters. Once all combinations are used to generate Moran simulations, the combination of parameters that provides the lowest error is taken to be the optimal fit to the data. The complete process for the first data fitting is shown in the algorithm in the Appendix. Here, values for the fitness were checked between 1 and 2 with a step size of 0.001.

In the case of fitting to treatment data, two datasets are provided so the fitness ratio *f*_*c*_/*f*_*h*_ is first fit to the vehicle data, and then the treatment efficacy is fit to the data depicting growth under treatment, using the determined fitness ratio. Here, other information about the data is estimated, such as the total number of cells, *N*, and the amount of cell divisions per hour. *N* is estimated by examining the growth shown in the data and visually extrapolating the total number of cells, and the cell divisions per hour are estimated to be *N*/8 as Ehrlich cells (those considered in the data) take eight hours to divide on average [50]. Simulations were run for treatment efficacies between 0% and 100% with a step size of 0.1%.

## 3. Results

### 3.1. Prediction for cancer evolution under two cell Moran model

To qualitatively confirm the Moran model’s ability to capture the growth of a population of cancerous cells, we simulate the model with no mutation (*r*_*m*_ = 0) starting with 10% cancer cells. A typical example of this is seen in **Figure 3A**, with the average and standard deviations shown in **Figure S1A**. When the fitness of both cell types is equal (*f*_*c*_ = *f*_*h*_ = 1), as expected, the percentage of type *c* cells remains around its initial value in the short term, however, for sufficiently long simulations, we expect one population to fixate given we have a finite population size, as seen in **Figure S1D** and by the expanding standard deviations in **Figure S1A**. If type *c* cells are given an advantage in the form of a higher fitness, i.e. *f*_*c*_ = 5 and *f*_*h*_ = 1, the percentage of cells increases logistically (**Figure 3B**), and there is much less deviation between simulations as seen in **Figure S1B**. It is expected that for any *f*_*c*_ > *f*_*h*_, the cancerous cell type *c* will reach fixation over time, but this will only be observed in simulations if the total number of cell divisions, *M*, is large enough. For example, fixing the number of cell divisions to *M* = 100 and varying the fitness, we see that the average cancer population does not reach fixation in this time, despite a 10-fold fitness advantage (**Figure 3C**), where this observation considers the average over *n*_*total*_ = 1000 simulations (see **Figure S1C** for the effect of the choice of *n*_*total*_ on the average simulation value).

We sought to understand further the influence of the total number of cell divisions on the ability of the cancer population to grow **(Figure S1E**). We see that the average cancer population only reaches fixation or a high proportion of the total cell population if the fitness value is high and the number of cell divisions is also sufficiently high. This suggests that cancer cell fitness can be quite low, and a cancer will still evolve if it is given sufficient time to take over the healthy cell population. We quantified the minimum fitness value needed for a given number of cell divisions so that the tumour population which reach over 50% (**Figure 3D**). We found there is a clear asymptote at around 67 cell divisions, which implies that it is impossible for cancerous cells to reach 50% given less than 67 cell divisions for this value of *N*. This is because the cancerous cells start with only one cell and are only able to increase by one cell at each time step, with a higher chance of death as the cancer population increases. As such, there must be some required time (in this case 0.5*N* or larger) before these cells can reach a threshold.

### 3.2. Evaluation of treatment efficacy under cancer cell mutation Moran model

Moran models are regularly used to track the probability of tunneling, which occurs when a cancer mutant reaches fixation before the original cancer [42], [51]. Komarova [42], Haeno *et al*. [43] and Jackson *et al*. [44] each consider a Moran model with three cell types, where the third type is a mutation from the original cancer with higher fitness. Each of these models discover that tunneling occurs naturally at some rate depending on the fitnesses of the cancerous and mutated cells. Using our model with 20,000 cell divisions, it is found that with our choice of parameters (*N* = 1000, *M* = 20,000, *f*_*h*_ = 1, *f*_*c*_ = 1.1, *f*_*m*_ = 1.5, *r*_*m*_ = 0.01) there is around a 0% chance, of tunneling, but if *f*_*m*_ = 3 then there is a 96.5% chance of tunneling. The average simulation under these conditions is in **Figure 4A** where the higher fitness is considered, also displaying one standard deviation away from the mean. To quantify the likelihood of tunneling given varying cancer cell and mutant fitnesses, we determine the proportion of simulations resulting in tunneling and have coloured the region of our (*f*_*c*_, *f*_*m*_) parameter space giving rise to tunneling or each cell type, seen in **Figure 4B**.

**Figure 4.**
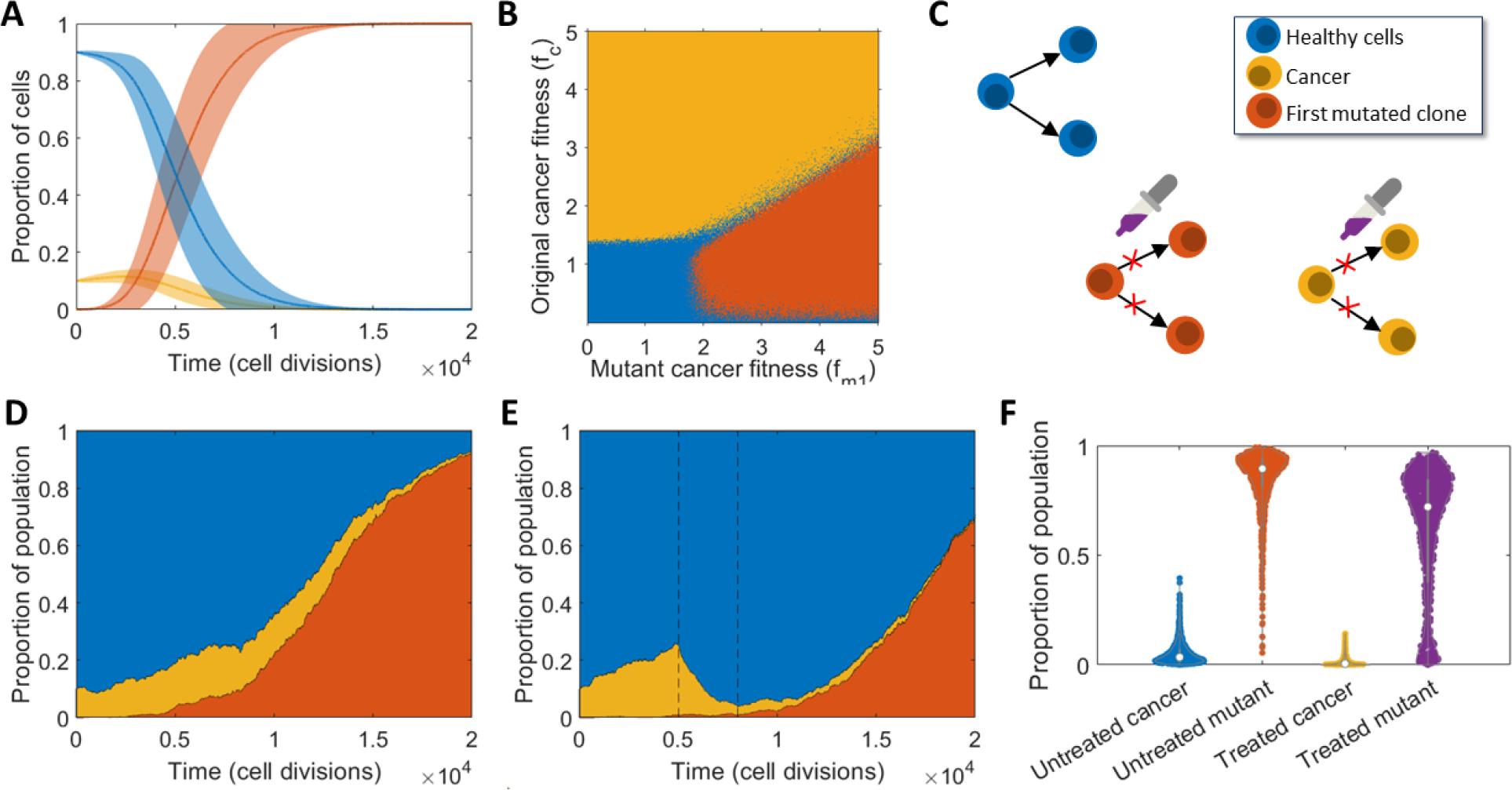
The impact of mutations on cancer growth and treatment. (A) Simulation of the Moran model showing tunneling of the mutant cancer, under high mutant fitness (*f*_*c*_ = 1.1*f*_*h*_, *f*_*m*_ = 3*f*_*h*_), average and standard deviation of *n*_*total*_ = 1000. (B) Regions depicting the most likely cancer type to fixate under varying *f*_*c*_ and *f*_*m*_, with the blue region representing little to no chance of either type fixating, orange representing a higher change for the original cancer type to fixate, and red representing a higher change for the mutant cell type to fixate. Further quantitative information is given in **Figure S2D**. (C) Model schematic depicting the effect of treatment in blocking or reducing the likelihood of cancer and mutant type proliferation. (D-E) Typical simulations depicting relative area of cell types over time with the inclusion of a fit mutant clone under (D) no treatment and (E) single high dosage treatment active for 3000 cell divisions. The period for which treatment is active is denoted by dashed black lines. (F) Comparison of the final proportions of both cancer types (cancer and mutant) without treatment and under single high dosage treatment for *n*_*total*_ = 1000 simulations. Presented as violin plots overlayed with individual data points.

We see there are clear regions of the (*f*_*c*_, *f*_*m*_) parameter space that give rise to one cell type outperforming the other. In particular for *f*_*m*_ > 2 and *f*_*c*_ > 1.5 there is a ratio of *f*_*c*_ /*f*_*m*_ ≈ 0.6, either side of which one cancer cell type wins. For example, *f*_*c*_ > 0.6*f*_*m*_ results in the original cancer outperforming the mutant. Largely, this image is also affected by the probability of mutation occurring *r*_*m*_ . For a further breakdown of the fixation proportion of each cell type see **Figure S2D**.

To investigate the impact of mutation on cancer growth under treatment, we simulated the cancer cell model with a single mutated clone. Using the choice of parameters described previously, a typical Moran process is simulated and displayed in **Figure 4D**, alongside a similar simulation with treatment applied in **Figure 4E**. Here, treatment begins at 5000 cell divisions and ends at 8000 cell divisions, with 80% efficacy against the original cancerous cells (type *c*) and 40% efficacy against mutant cells (type *m*). We find that the mutant clone will take over even in the presence of treatment if it is present when treatment begins. Given that the conditions are such that tunneling of the mutant clone is expected to occur, it is also expected that treatment may fail to control the mutant clone. As a result, the tumour’s response to the treatment may also be examined to determine if tunneling is occurring with a resistance cancer clone. Further examples of these behaviours may be seen in **Figures S2A** and **S2C**. Using these parameters, the final proportions of cancer types *c* and *m* were recorded and examined with and without treatment as seen in **Figure 4F**. Here it can be seen that treatment reduces the final proportions of each cancer type, though there remains a high possibility that the mutant clone approaches fixation with large variation in the final mutant number, suggesting that in some instances, the healthy cells may regain control.

### 3.3. Evaluation of treatment strategies

Using the Moran model with three cell types (healthy *h*, cancerous *c*, and mutant *m*), different treatment strategies were tested and compared (**Figure 5A**): applying the maximum tolerated dose (MTD) after mutant growths arise, earlier dosing, dual dosing, sustained dosing, and waning treatment. Here, the MTD is assumed for all simulations except the sustained dosage and waning treatment. The sustained dosage is a reduced dosage (lower treatment efficacy) that is applied for a longer period, and the waning treatment is an exponential decrease of the effect of treatment to capture pharmacokinetic drug dynamics. The dual dosing treatment considered that the mutant cancer was targeted in the first dose and that the second dose targeted the original cancer. See **Figure 5B-5F** for the mean and standard deviation of 1000 model simulations.

**Figure 5.**
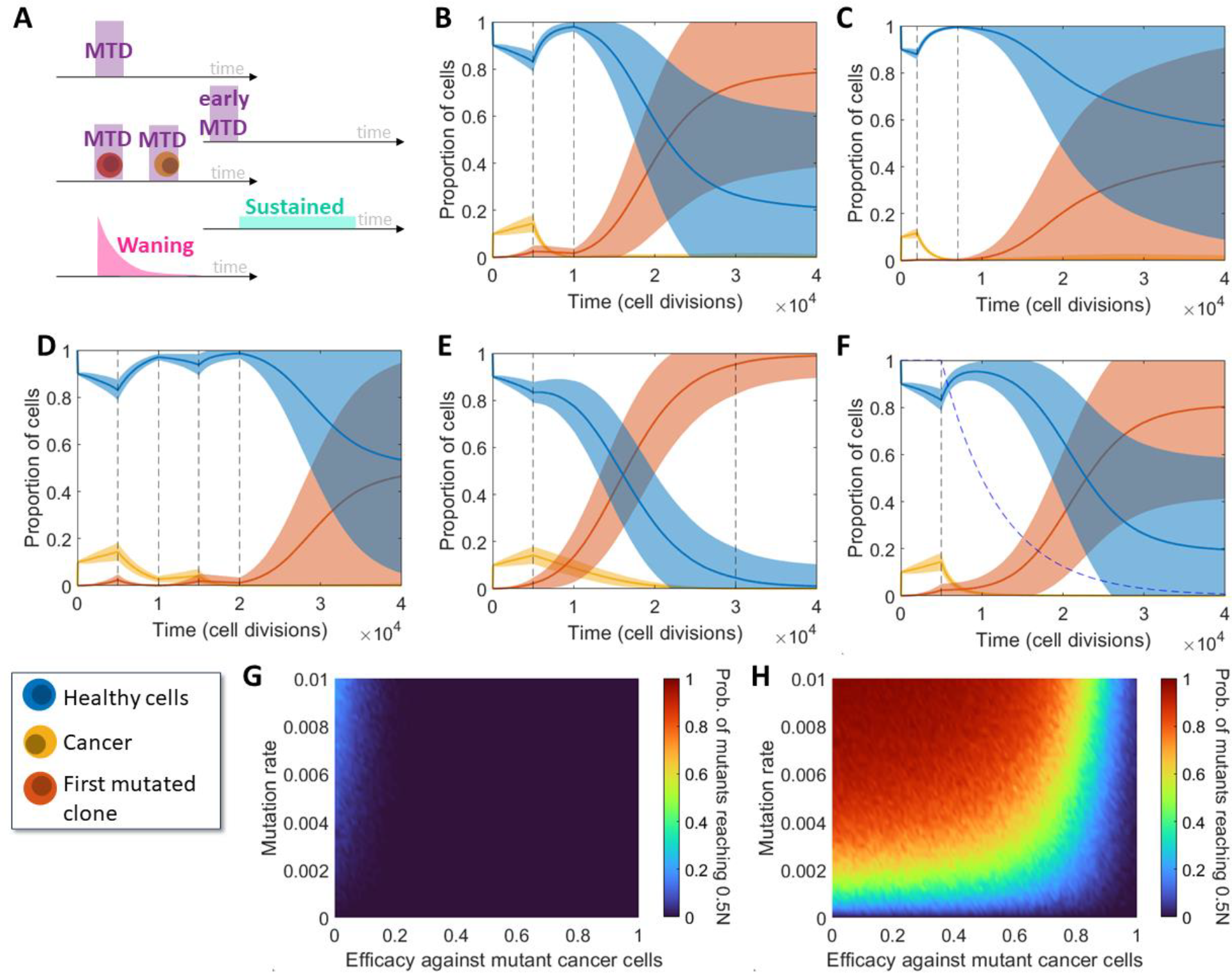
Comparisons between different treatment methods and models. (A) Descriptions of the five treatment methods considered in (B-F): a MTD, an early MTD, a dual dosage (one targeting the mutant followed by one targeting the cancer), a sustained low dose and a waning treatment. (B-F) Simulations of cancer growth under varying treatment strategies, average and standard deviation of *n*_*total*_ = 1000. (B, C) Single high dosages applied for *t*_*length*_ = 5000 cell divisions between the vertical dashed lines (B) after the mutant clone is expected to arise and (C) before the mutant clone is expected to arise, i.e. early treatment. (D) Dual dosing with the first targeting the mutant cancer clone and the second targeting the original cancer clone. (E) Sustained dosing with an efficacy one fifth of that in other tests. (F) High dosage with a waning drug concentration modelled by the blue dashed line. (G, H) Proportion of simulations (here *f*_*m*_ = 1.6 and *M* = 20,000) with varying *r*_*m*_ and eff_*m*_ where mutant cells reach a population of 0.5*N* under (G) single high dosage treatment, and (H) sustained treatment with efficacy scaled by 1/3 and length scaled by 5. Corresponding two parameter plots for the original cancer are given in **Figure S2B** and **S2E**.

In general, the presence of a mutated cancer clone provides the cancer growth with an inherent treatment resistance which makes treatment difficult. In certain cases, particularly the MTD, sustained treatment and waning treatment, the presence of the drug only aided the growth of the mutated clone by reducing the original cancer clone. As such, applying the MTD can be effective assuming that the treatment is effective towards the mutated clone or if the treatment dose is high enough, but can fail to eliminate the mutated cancer otherwise (see **Figure 5B**). Sustained treatment often fails similarly since the reduced dosage is not strong enough to eliminate the mutated cancer (see **Figure 5E**).

This motivates the use of early dosing which tends to eliminate the cancer before any mutation arises and does not require a large dosage if targeted towards the original cancer, as seen in **Figure 5C**. Finally, dual dosing was considered to target the mutated cancer in a first dosage to reduce or eliminate it, and then apply a second dose targeted towards the original clone. This provided varying results depending on mutation (see **Figure 5D**), but it is again determined that intra-clonal heterogeneity presents challenging resistances to treatment.

An alternative treatment model considering a waning effect on the drug’s concentration is considered in **Figure 5F**, where the concentration is modelled by a decaying exponential based on the drug’s half-life. In this case, the half-life is arbitrarily selected to be 5000 cell divisions, though this may be more appropriately selected for specific cases. Here, the waning treatment model represents a more natural transition between active and ineffective treatment, in this case showing results less promising than consistently administering the MTD, and more promising than administering a consistent sustained dose. Individual model trajectories for the five treatment scenarios are plotted in **Figure S3** and corresponding violin plots for the final proportion of cancer and mutant cells is given in **Figure S4**.

**Figures 5G** and **5H** compare the mutant cancer (type *m*) responses to treatment with the MTD and sustained treatment respectively, where the mutation rate and efficacy against mutant cells vary. These plots consider the proportion of simulations where mutant cells grow to at least 50% of the total cell population *N* within *M* = 20,000 cell divisions. Additionally in **Figures S2B** and **S2E**, a similar comparison is made for the original cancer cell type (type *c*). For the selected parameters, the original cancer (cell type *c*) will almost never reach 50% of *N*, which is likely due to tunneling rather than treatment, as the mutated cancer is expected to take over quickly. **Figures 5G** and **5H** show that the mutated cancer cell clone (type *m*) is more likely to grow to a substantial size in the sustained treatment scenario if the mutation rate is high and treatment is not as effective against that cell type, as expected. Specifically using the MTD in this case, low efficacy is required for the mutated cancer to have a non-zero chance of reaching 50% of *N*. It is evident that treatment with the MTD is overall more effective than sustained treatment when changes in efficacy and mutation rate are considered. Some selections of these parameters can facilitate effective sustained treatment such as zero mutation rate or a treatment efficacy of 100%, however these selections are not practical.

### 3.4. Herceptin case study

To validate the proposed model, the recursive form in Eq. (5) is fitted to the same data used by West *et al* [41], originally published by Laird [48], [49]. Two parameters are fit to the data: the ratio of cancer fitness to healthy cell fitness, *f*_*c*_ /*f*_*h*_ and the number of cell divisions per unit on the horizontal axis, assuming that there is one initial cancer cell. The Root Mean Square Error (RMSE) between the model fit and the data for different cancer fitnesses is confirmed to be a minimisation of the RMSE, as seen in **Figure S5A**. The recursive model is used rather than an average of simulations as it provides a much faster runtime when comparing results from many combinations of parameters. It is found that a fitness ratio of *f*_*c*_ /*f*_*h*_ = 1.1 most accurately represents the data with 8998 cell divisions corresponding to one unit along the horizontal axis. **Figure 6A** shows that the model fitted using the recursive model proposed in this work fits the raw data closely. The recursive model involving mutations to type *m* cells was fit to the same data set (results not shown). This involved fitting both fitness ratios *f*_*c*_ /*f*_*h*_ and *f*_*m*_/*f*_*h*_, the mutation rate *r*_*m*_, and the number of cell divisions per unit at the same time, assuming that the data begins with no mutated cells. Interestingly, the best fit was found when *r*_*m*_ = 0, corresponding to the model fit with only one cancer clone, suggesting that the cancer described in the data is not genetically complex.

**Figure 6.**
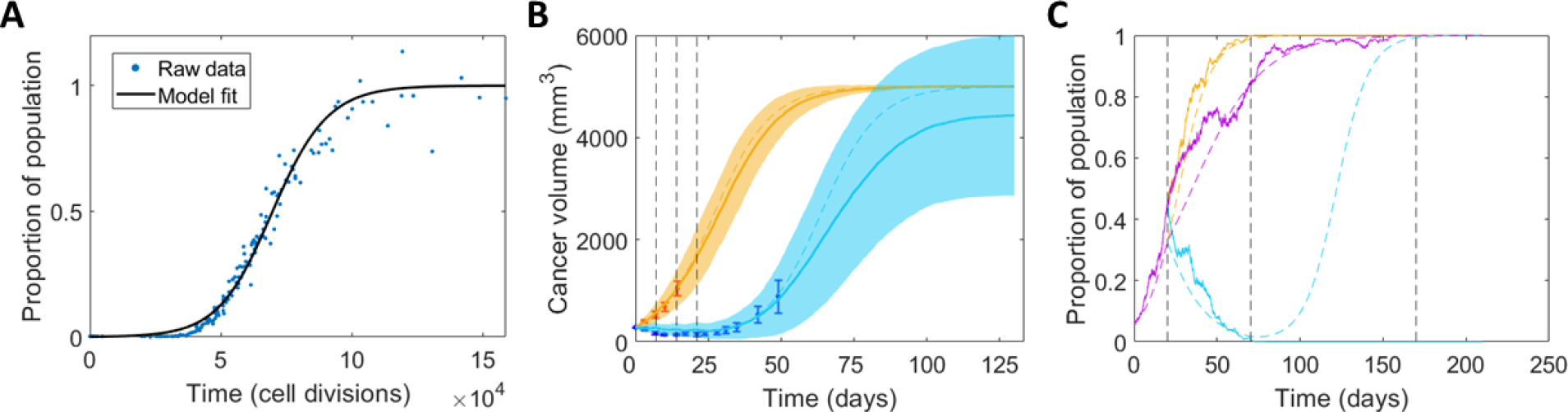
Summary of model fitting to experimental data and testing. (A) Comparison between data depicting murine mammary cancer growth [48], [49] and the recursive model fit to the data. (B) Comparison between data and model fits for vehicle cancer growth data (orange) and growth under treatments applied four times (blue) [12] starting at zero days and re-administered every seven days with waning drug concentration, simulations shown with mean and standard deviation over *n*_*total*_ = 100. (C) Comparison between treatment techniques with waning drug effects using the model fit to the data in (B), with a single high dosage starting at 20 days and lasting for 50 days (period within first two black dashed lines), and a sustained dosage starting at 20 days with efficacy scaled by 1/3 and length scaled by 3.

Similarly, this model can be used to provide a framework for testing treatments against Ehrlich ascites carcinoma. Given that Ehrlich cells divide around every eight hours [50], it can be assumed for this model that the amount of time per simulated cell division is 8/*N*. Additionally, this cancer may be treated with Herceptin, which is found to have an average half-life of 5.8 days [52], [53]. Using this information, the model could then be fit to other data measuring the growth of HER2-Positive breast cancer tumours in mice with and without Herceptin treatment presented by Lewis Phillips *et al*. [12]. **Figure 6B** shows untreated cancer growth data against a fitted model, following the same process as before. This finds that the cancer considered has a fitness of around *f*_*c*_ = 1.035*f*_*h*_. This information is then used alongside the information about the half-life of Herceptin and the dosages indicated by Lewis Phillips *et al*. [12] to fit the recursive model to data with treatment, assuming a waning effect (Eq. (4)), also seen in **Figure 6B**. It is found here that this treatment is around 5.5% effective against this cancer growth (i.e. eff_*c*_ = 0.055). It is seen in **Figures S5B** and **S5C** that each of these parameters minimise the RMSE between the data and the model, respectively.

We then considered these parameters in our previous treatment scenarios from **Figure 5A**, however, particularly focussing on MTD. Results shown in **Figure 6C** compare a single high-dose injection (MTD) to a sustained low-dose injection, both with waning drug concentration after treatment concludes. It is seen from the given typical simulations that this sustained treatment with Herceptin does not have a strong impact on the cancer’s growth, though the single high-dose treatment is able to eradicate the cancer entirely. Alternatively, **Figure S5D** considers fitting the model to the same data assuming single high dose injections without waning treatment, which suggests a treatment efficacy of 4.3%. Using these fit parameters, the expected outcome from treatment methods such as applying a higher dosage and sustained treatment can be simulated as seen in **Figure S5E**. Similarly, it is found that sustained treatment gains no advantage over high dose injections and does not cause death to as many cancer cells.

## 4. Discussion

Moran models present a useful mathematical tool that can be used to capture the occurrence of resistant-mutant clones in the cancer population and the impact of this has on cancer therapy. Resistance is one of the major causes of cancer related deaths, and understanding how the presence of resistance cancer subpopulations impact new therapeutic designs is crucial to the drug development pipeline. In this work, we develop a simple stochastic model for cancer proliferation and mutation using a Moran Process. With this model, we investigate the impact of therapeutic design on treatment efficacy focusing on two main treatment protocols: short treatment windows with the MTD administered and longer treatment windows with lower treatment dosages. These are referred to as MTD and sustained treatment. We then examined how a model such as this could be used to match to experimental data.

Using a simple two cell (cancer and healthy cell) Moran model, we replicated the biological behaviour expected of a non-cancerous and cancerous subpopulation, whereby, cells with a fitness advantage grow logistically. Most interestingly, we quantified the minimum fitness needed for a given number of cell divisions to result in the cancer growing to more than 50% of the population. We found that for a fixed number of cells (i.e. 100 cells), as the number of cell divisions increased, the minimum fitness needed to grow the tumour decreases exponentially. There was a clear asymptote providing the minimum number of cell divisions needed for the tumour to grow beyond 50% of the population. This asymptote is intuitive and represents the minimum number of timesteps needed to increase the tumour population from 1 to 50. Introducing mutations, we were then able to quantify the regions of the mutation parameter space that would provide successful tunnelling for the mutant population, i.e. *f*_*c*_/*f*_*m*_ < 0.6.

Investigating the effects of a MTD treatment, simulations first revealed the large variation in responses to treatment of the mutant population. For some trajectories, there were no mutants and minimum cancer cells, suggesting the healthy cell population had regained control. Furthermore, the final average mutant population is decreased in the presence of treatment. Comparing the MTD to sustained treatment, it was clear that MTD is more likely to be effective than the sustained treatment for a range of cell fitness, i.e. cancer growth rates. This is an important finding given the interest in using sustained dosages and suggests that more work is needed to investigate the impact resistance may have on this treatment. We successfully calibrated the Moran model to *in vivo* control and breast cancer treatment data using our recursive form of the Moran model. We found that in this parameter regime our results were corroborated with MTD being most effective and sustained treatment being virtually ineffective.

We found that early treatment is more effective than later treatment and this is unsurprisingly well supported in the literature as treatment should be administered early in the cancer’s growth to reduce the effect of the evolutionary process of cancer mutation and reduce the tumour’s genetic complexity. This result was also found by West *et al*. [39] who suggested that treatment should be administered in the early stages to prevent genetic complexity. In addition to early treatment, we also found dual dosing of treatment was more effective than other treatments, such as waning treatment. This suggests that targeting the more fit mutant clone may be more beneficial, though large variation was seen in our results. In the past decade, multiple clinical trials have investigated the use of intermittent therapy, which applied on/off treatment cycles after an initial induction period [54]. These therapies exploit the assumption that intermittent treatment strategies exploit evolutionary principles to delay the onset of resistance while decreasing accumulated drug doses and reducing toxicity [54]. These therapies have mixed results [54].

Waning treatment was found to be just as ineffective as sustained treatment, suggesting that the waning of treatment efficacy, due to pharmacokinetics might be what causes issues with efficacy. Furthermore, this suggests that the waning treatment (if more representative of true human dynamics) should be what we compare to sustained treatment and comparing the violin plots for the final amount we see there is more variation in the sustained treatment then the waning treatment – suggesting some people might response positively to sustained treatment.

There are limitations to our currently modelling framework and future work can look to improve and also further validate experimentally the predictions of the model. For example, we didn’t consider a cost associated with resistance to therapy, and it has been suggested that if sensitive population are much larger than the resistant population in a cancer, it usually indicates that there is a high fitness cost for resistance [54]. Fortunately, the ratio of sensitive-to-resistant cells can often be inferred from the initial response to therapy [54] and future work hopes to use informed initialisation of the model. In addition, we are using a non-spatial model of cancer cell behaviour to capture what is often a very spatial process. Given that our model is sufficiently simple and does not try to link proliferation and death to real tumour processes, it can be thought to capture a range of biological phenomena occurring in the tumour.

In this work, we present a simple Moran framework that can be used to quantify the impact of resistance on a range of treatment modalities. Most significantly, we find that early treatment is the most effective form of treatment, followed closely by dual treatment where the mutant clone and then the cancer clone are targeted in succession. MTD is an effective therapy, when compared to sustained dosages and this is most seen most clearly in the case where the model was matched to experiment measurement for breast cancer treatment with Herceptin. This study suggests more work needs to be done to investigate sustained dosage therapies in clonal heterogeneous subpopulations to investigate whether resistance is going to hinder this treatment’s effectiveness.

## Acknowledgements

ALJ acknowledges support from the QUT Women in STEM Writing Retreat and funding from the ARC Discovery Grant DP230100025.

## 5. Appendix

### 5.1. Algorithm for the first data fitting

#### Algorithm 1 Fittig *M*/*t* and *f*_*c*_*/f*_*h*_ to vehicle data

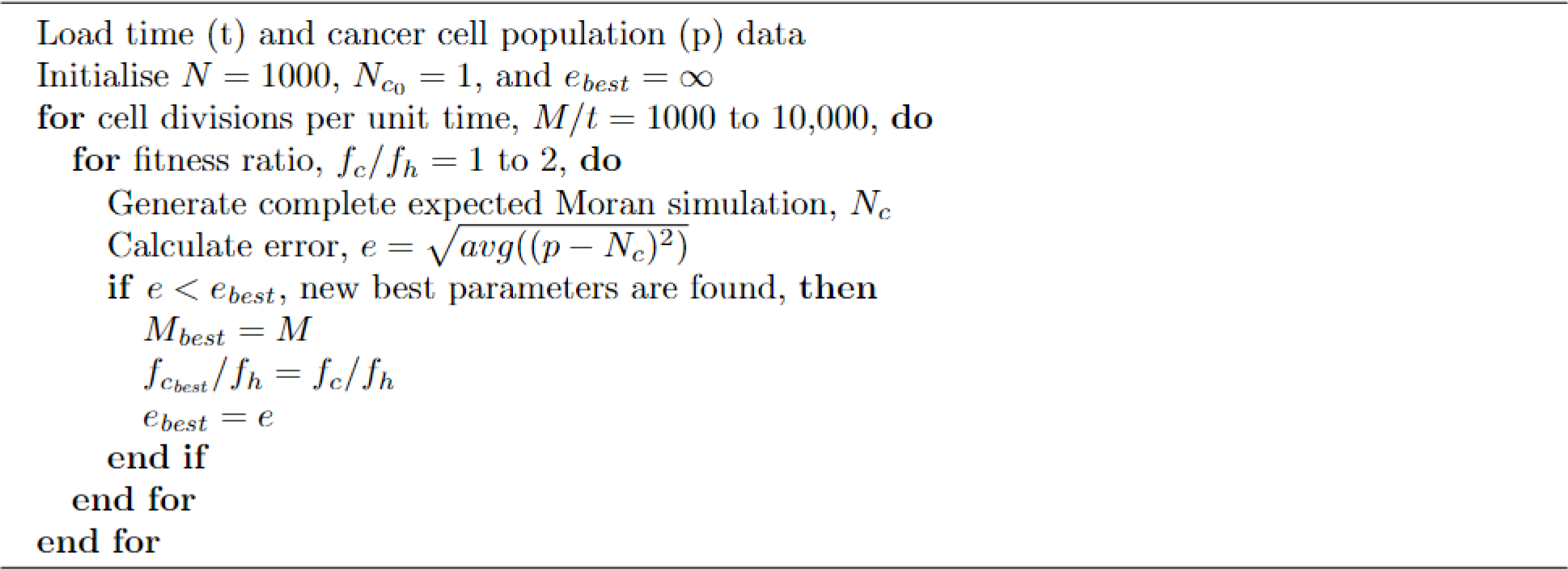

### 5.2. Full list of model transition probabilities: healthy cells, cancer cells, mutant cancer cells and treatment

The probability of a cell of type *h, c* or *m* dying respectively is given by:

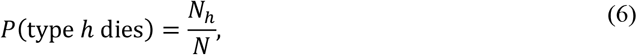

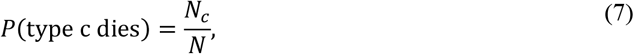

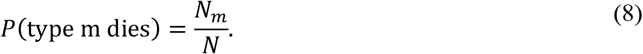

The probabilities of cells of type *h, c* or *m* reproducing under the effects of treatment (see Eq. (3)-(4)) are:

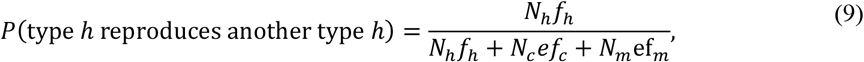

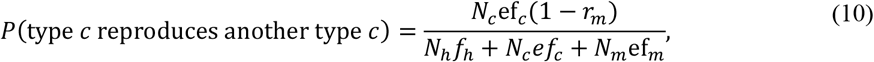

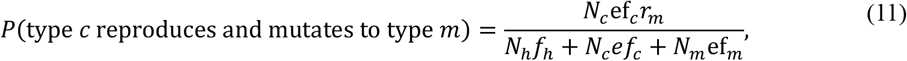

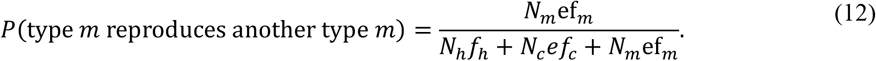

### 5.3. Supplementary figures and tables

**Figure S1.**
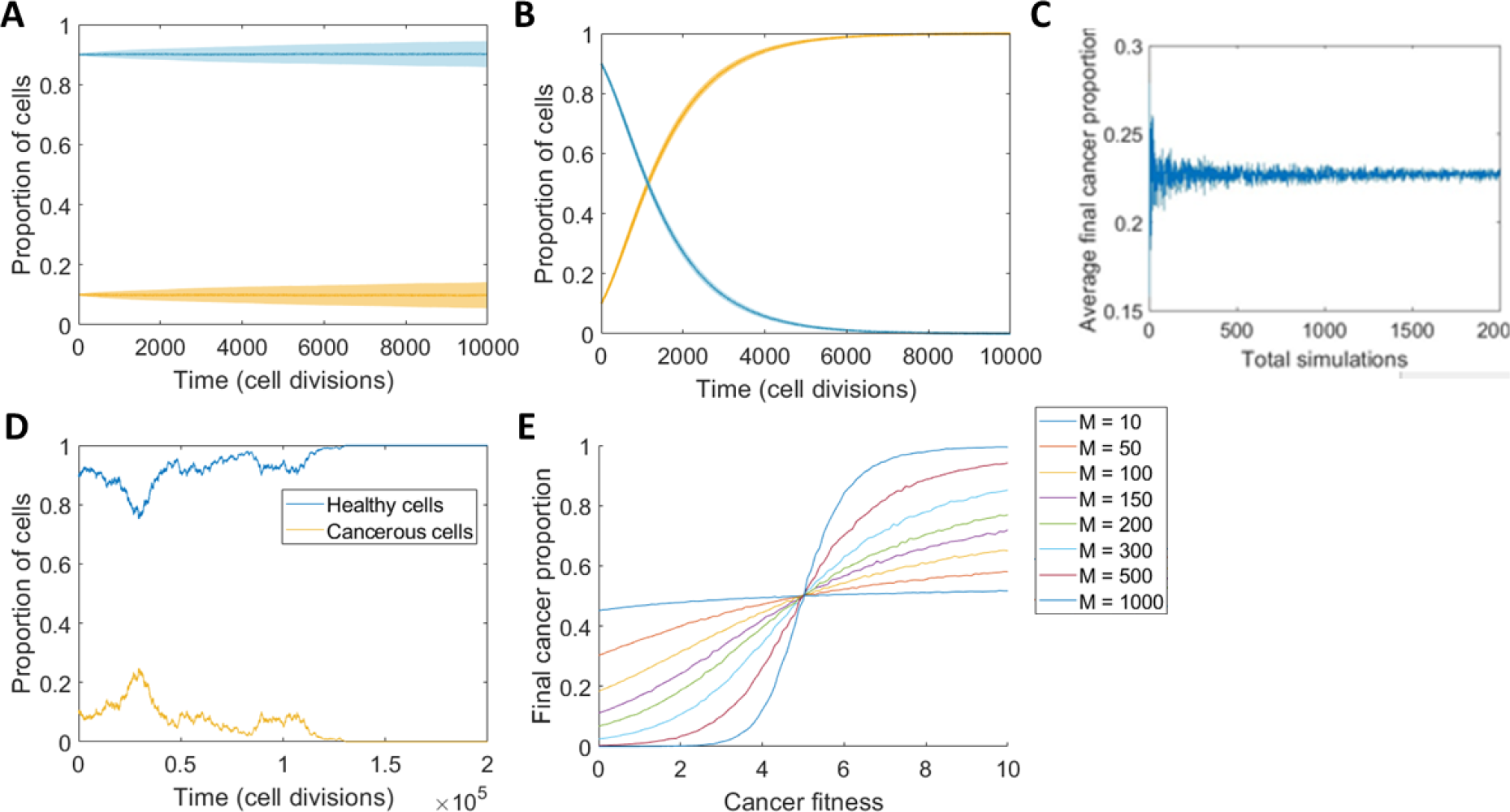
Moran model outputs for cancer growth in the absence of mutant clones and treatment. (A, B) Simulations of the Moran model for (A) equal fitness (*f*_*c*_ = *f*_*h*_ = 1) and (B) unequal fitness (*f*_*c*_ = 5, *f*_*h*_ = 1), with mean and standard deviation for *n*_*total*_ = 1000 and an initial proportion of cancer cells (yellow) or 10% and healthy cells (blue) 90%. (C) To assess the convergence of the stochastic process the average final proportion of cancer cells under a typical simulation is given as *n*_*total*_ increases. (D) Typical simulation over large time for equal fitness (*f*_*c*_ = *f*_*h*_ = 1), i.e. corresponding to (A) and (B). (E) Comparison between the final proportion of cancer cells as fitness varies for *M* ∈ [10,1000], where *M* is the total number of cell divisions.

**Figure S2.**
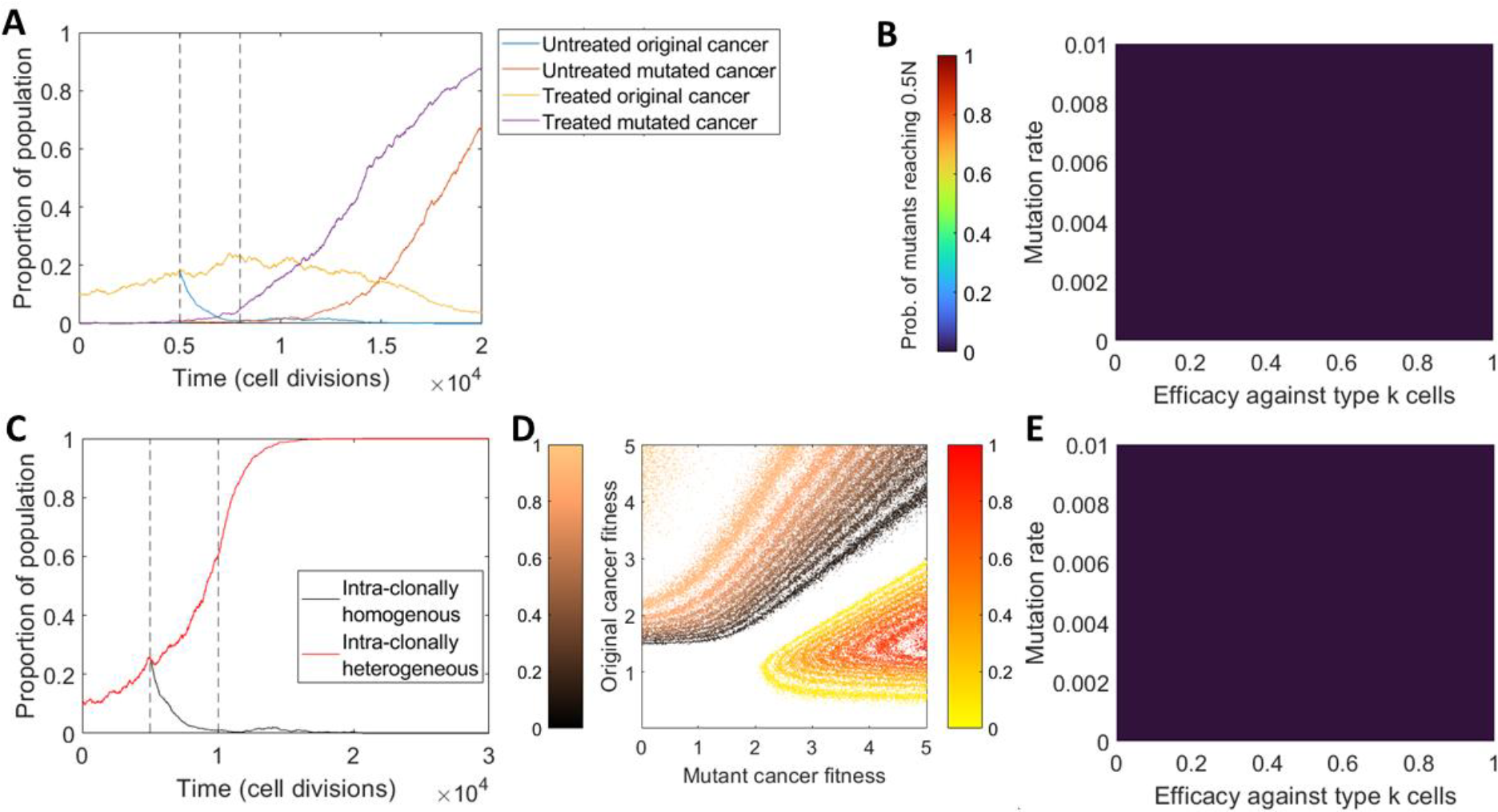
Supplementary figures summarising the impacts of mutant clones on cancer growth and treatment. (A) Typical simulation of intra-clonal heterogeneous cancer growth under single high dosage treatment. (C) Comparison between the effects of treatment on a cancer growth with and without the presence of a mutant clone, given typical simulations. (D) Proportion of *n*_*total*_ = 100 simulations in which each cancer type fixates as *f*_*c*_ and *f*_*m*_ vary. (B, E) Proportion of simulations with varying *r*_*m*_ and eff_*m*_ where original cancer cells reach a population of 0.5*N* under (B) single high dosage treatment, and (E) sustained treatment with efficacy scaled by 1/3 and length scaled by 5.

**Figure S3.**
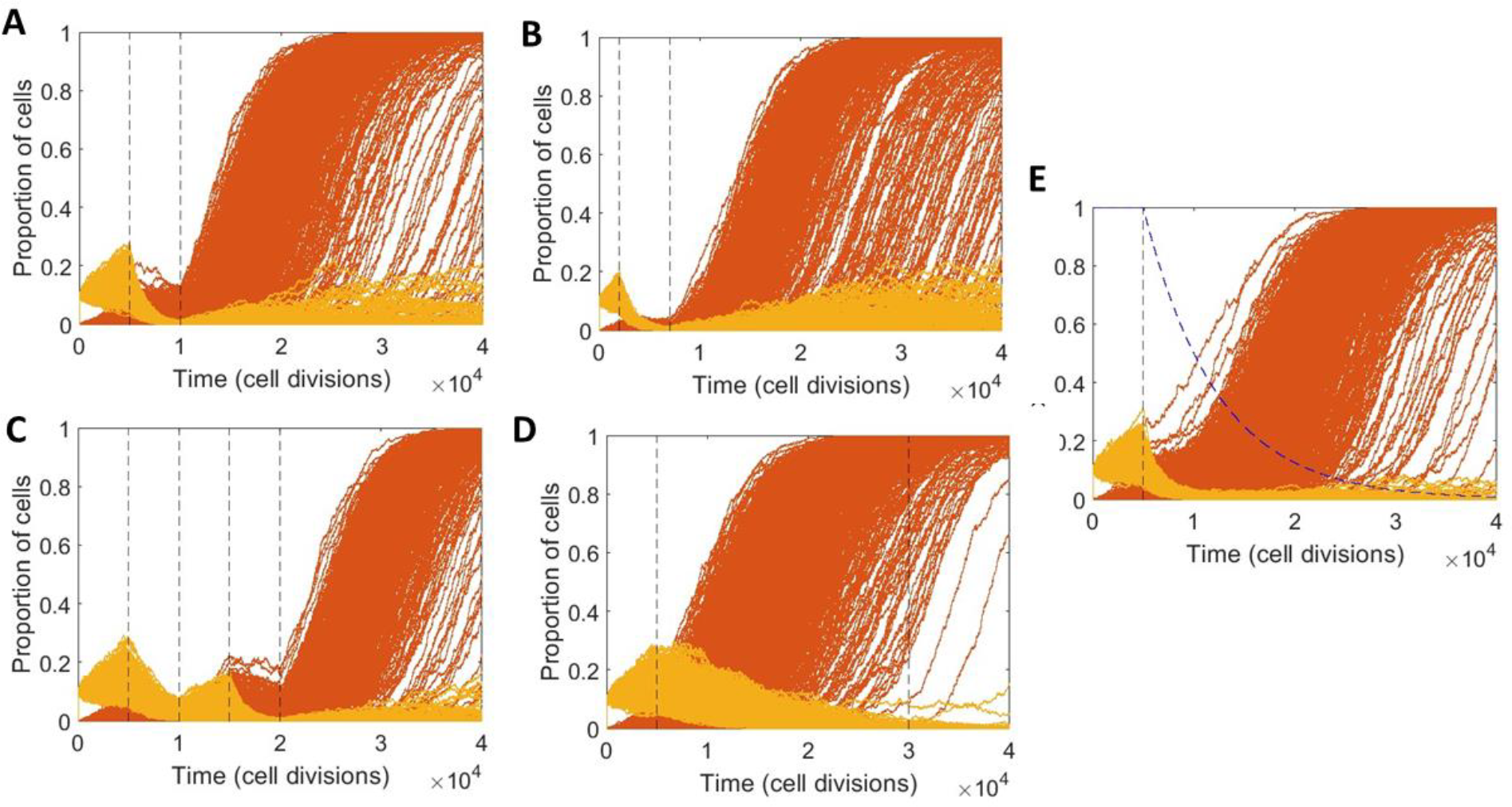
Supplementary figures for comparisons between simulations of different treatment methods and models. Plots of *n*_*total*_ = 1000 individual simulations of original and mutant cancer growths under treatment. (A, B) Single high dosages applied for 5000 cell divisions between the vertical dashed lines (A) after the mutant clone is expected to arise and (B) before the mutant clone is expected to arise. (C) Dual dosing with the first targeting the mutant cancer clone and the second targeting the original cancer clone. (D) Sustained dosing with an efficacy one fifth of that in other tests, applied over 25,000 cell divisions. (E) High dosage with a waning drug concentration modelled by the blue dashed line.

**Figure S4.**
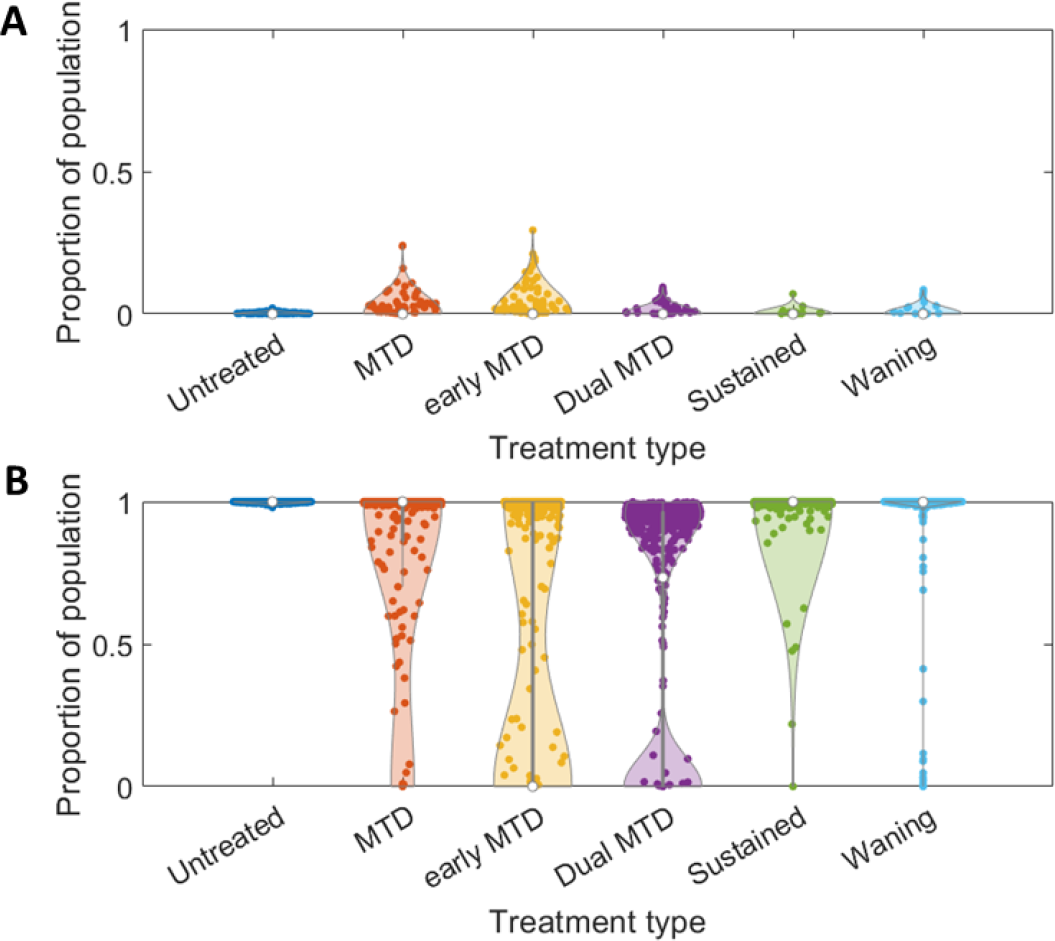
Supplementary figures for comparisons between final proportions of cancer under of different treatment. Final proportions of cancer growths for *n*_*total*_ = 1000 simulations under treatments including respectively: single. high dosages applied for 5000 cell divisions after the mutant is expected to arise and then before it is expected to arise, dual dosing with the first targeting the mutant cancer clone and the second targeting the original cancer clone, sustained dosing with an efficacy one fifth of that in other tests, applied over 25,000 cell divisions, and high dosage with a waning drug concentration modelled by the blue dashed line seen in **Figure S4E**. These treatment methods are tested simulations involving both (A) the original cancer clone, and (B) the mutant cancer clone.

**Figure S5.**
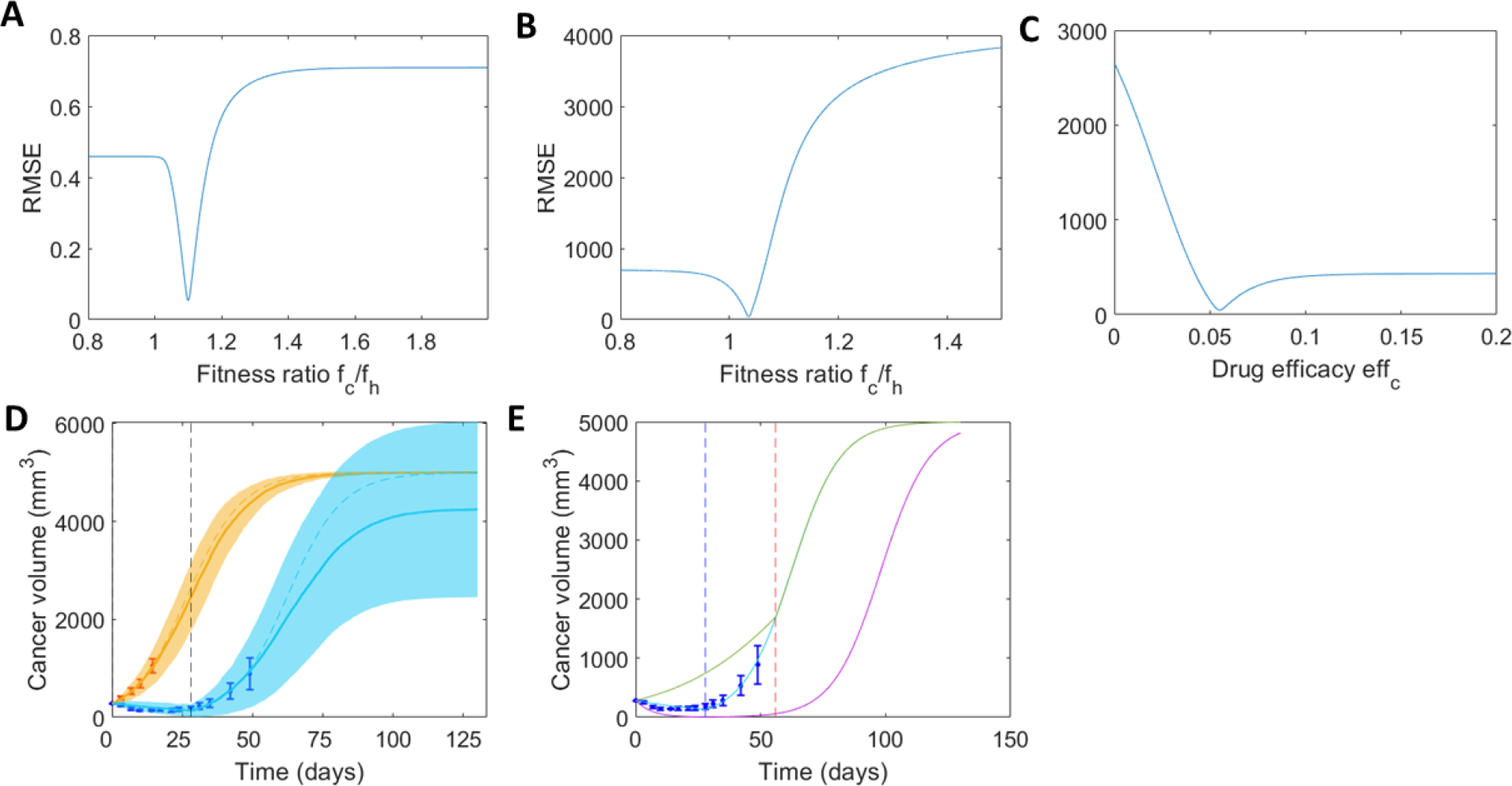
Supplementary figures summarising model fitting parameters and alternating treatment fitting. (A-C) RMSE between the recursive model and data for different parameters. (A) Fitness ratio *f*_*c*_/*f*_*h*_ fit to murine mammary cancer data [48], [49]. (B) Fitness ratio *f*_*c*_/*f*_*h*_ fit to murine mammary cancer data without treatment [12]. (C) Drug efficacy eff_*c*_ fit to cancer data under treatment [12], using the optimal fitness ratio observed from (B). (D) Alternate comparison between data and model fits for vehicle cancer growth data (orange) and growth under consistent treatment over four weeks (blue) [12] without waning drug concentration, simulations shown with mean and standard deviation over *n*_*total*_ = 100. (E) Comparison between expected cancer growth under the treatment suggested in the data, treatment with a higher dosage, and sustained treatment with efficacy scaled by 1/2 and length scaled by 2.

**Table 1.** List of model variables and parameters. Below is a summary of all parameters/variables in the model. Relevant references for all the parameters is given.

| Symbol | Description | Range |
| --- | --- | --- |
| $N$ | Total number of cells | [100,10000] |
| $M$ | Total number of cell divisions | [100,100000] |
| $f_c$ | Fitness of cancer cells (also applicable to all other cell types) | (1,2] |
| $N_{c_n}$ | Number of cancer cells at timestep $n$ (also applicable to all other cell types) | [0,$N$] |
| $r_{m1}$ | Mutation rate for type $m1$ mutant clones (also applicable to other cancer mutants) | [0,0.01] |
| $eff_c$ | Efficacy of treatment against cancer cells (also applicable to cancer mutants) | [0.4,0.8] |
| $w$ | Weakness in treatment efficacy compared to a single high dose | [2,5] |
| $t_{start}$ | Starting time for treatment (in cell divisions) | [0,5000] |
| $t_{length}$ | Length of treatment (in cell divisions) | [2000,5000] |

